# Long-read sequencing reveals extensive size variation and suggests evolutionary patterns of expansion in bivalve mitogenomes

**DOI:** 10.1101/2024.12.23.625590

**Authors:** Ning Zhang, Lu Qi, Biyang Xu, Lingfeng Kong, Qi Li

**Affiliations:** Key Laboratory of Mariculture, Ministry of Education, Ocean University of China, Qingdao, China; Laboratory for Marine Fisheries Science and Food Production Processes, Qingdao National Laboratory for Marine Science and Technology, Qingdao, China; South China Sea Fisheries Research Institute, Chinese Academy of Fishery Sciences, Guangzhou 510300, China; Institute of Marine Science and Technology, Shandong University, Qingdao 266237, China

**Keywords:** Mitochondrial genome size, Bivalve, Long reads, Assembly, Unassigned region

## Abstract

Bilaterian mitochondrial genomes are generally of conserved size and gene content, typically ranging from 14 to 20 kb and coding 37 genes. However, the major exception to this rule has been found in many bivalve species which allow the presence of long unassigned region (regions that are functionally unassigned). It is still unclear whether there are universal patterns to expansions across bivalves. Additionally, the prevalence of highly repetitive sequences within complex mitogenomes has prevented its complete assembly and led to its systematic omission from mitogenomic analyses. Here, leveraging the growing genomic resources in Mollusca, we devised a mitogenome assembly pipeline using long-read sequencing to characterize mitogenome features in 178 mollusk species. Comparing our assemblies with reference mitogenomes based on Sanger/short-read sequencing, we identified missing sequences, repeats and novel gene duplications and found that bivalves exhibit an exceptional diversity in mitogenome size compared to other mollusks. Subsequently, we investigated the factors potentially influencing mitogenome size, the potential functional elements and the patterns of expansion regions. Furthermore, we discussed that the possible selective pressure on mitogenome size and hypothesized that the variability of bivalve mitogenomes may be linked to innate immunity. Our findings underscore the significance of long-read sequencing in elucidating complex mitogenomes and provide new insights into the evolution of bivalve mitogenomes.

## Introduction

Mitochondria, specialized organelles possessing their own genome (mitochondrial genome or mitogenome), play a variety of crucial roles within the cell, far beyond their essential energetic functions in oxidative phosphorylation, the Krebs cycle, and fatty acid oxidation (Karnkowska, et al. 2016; Monzel, et al. 2023). Mitogenomes are the remainder of the genome of an α-proteobacteria ancestor, incorporated into the eukaryotic cell by endosymbiosis (Roger, et al. 2017). In eukaryotic lineages, mitogenomes display marked diversity in sequence size, ranging from 6 kb (malaria parasite) to >11 Mb (catchfly plant) (Sharma, et al. 2001; Sloan, et al. 2012). Compared to plants, fungi and numerous protists, most animal mitogenomes, particularly those of bilaterians are remarkably compact (14–20 kb), coding for 13 intron-less protein coding genes that belong to different enzyme complexes of the oxidative phosphorylation system, 2 rRNA (12S and 16S) and 22 tRNAs (Boore 1999; Gissi, et al. 2008; Lavrov and Pett 2016). However, with the development of sequencing technology, a broader sampling of bilaterian animal mitogenomes has found many cases that deviate dramatically from traditional description and vary widely in genome size, and Bivalvia (Mollusca) is especially replete with these exceptions (Ghiselli, Gomes-dos-Santos, et al. 2021).

Bivalves are an ancient and ubiquitous group of molluscs with over 10,000 described species in marine and freshwater environments (Ghiselli, Iannello, et al. 2021). They are economically significant as a food source (e.g. oysters, scallops, clams, cockles and mussels), and ecologically important in view of their filter-feeding habits and biomass (Gage and Tyler 1991; Pawiro 2010). Thanks to their remarkable diversity and peculiar biological features, they have been proposed as promising model organisms for investigating a broad spectrum of biological, ecological, and evolutionary issues, particularly in fields like mitochondrial biology and its evolution (Nicolini, et al. 2023). Bivalves harbor some of the most complex mitogenomes among metazoans, characterized by considerable size variation, significant rearrangements, gene duplications and losses as well as documented instances of doubly uniparental inheritance (Ghiselli, et al. 2013; Ghiselli, Gomes-dos-Santos, et al. 2021). For example, more than 13 mitogenomes of ark shells (Arcidae) have length larger than 28 kb, which display a markedly increased size and show a lineage-specific expansion (Kong, et al. 2020; Zhang, et al. 2022). The zebra mussel (*Dreissena polymorpha*) mitogenome is 67,195 bp, which is 3–4 times larger than that of most bilaterians (McCartney, et al. 2022). Moreover, large mitogenomes are also found in other bivalve lineages such as sea scallop *Placopecten magellanicus* (31-41 kb) (Smith and Snyder 2007) and freshwater pearl oyster *Pinctada imbricata* (31 kb) (Zhan, et al. 2018). These studies suggest that changes in mitogenome size are a common feature across many bivalve species, yet the mechanisms driving these alterations remain poorly understood, impeding our understanding of the relationship between particular mitogenomic characters and evolutionary fitness.

Changes in mitogenome size can result from a variety of factors, including gene duplication, gene elongation, and the expansion of unassigned region (regions that are functionally unassigned) (Lubośny, et al. 2022; Zhang, et al. 2022). Expansion of UR (unassigned region) is usually considered a significant force driving mitogenome size changes among these factors. Typically, URs have usually been considered short and comprise no more than 1.5 kb of the mitogenome (Lunt, et al. 1998). However, more recent studies on bivalves (Kinkar, et al. 2021; McCartney, et al. 2022) had revealed the presence of long and complex tandem-repetitive elements in URs of large mitogenomes, which pose a serious challenge to previous sequencing technologies and assembly efforts. Almost all mitogenome assemblies have traditionally depended on bioinformatic assembly of sequencing reads generated via Sanger sequencing or next-generation sequencing (NGS). Long-range PCR and Sanger sequencing relies on assumed gene order similarity with reference sequences. This approach faces challenges when significant gene rearrangements occur, often resulting in failed reactions, multiple amplicons, or unanticipated lengths (Bensasson, et al. 2001; Miller, et al. 2004). NGS has improved mitogenome assembly by sequencing multiple small DNA segments in parallel, allowing sequencing without prior knowledge of the sequence or gene order (Gan, et al. 2014). However, a major limitation of NGS is its difficulty in spanning complex regions, particularly repetitive regions and segmental duplications (Formenti, et al. 2021). Taken together, assemblies using both scenarios (Sanger or short-read sequencing) may give a uncomplete mitogenome that precludes the accurate inference of bivalve mitogenome structure, ultimately impacting evolutionary analysis.

In recent years, third-generation technologies, based on single-molecule and long-read sequencing, have been successful in resolving repetitive and structurally complex DNA elements (Dijk, et al. 2023). Long-read sequencing can span easily intricate regions to solve the overlap uncertainty issue of Sanger-based approaches and short-read NGS and uncover previously undetectable mitogenomic architectural features (Giani, et al. 2020). Several studies combining accurate Illumina reads with long reads to improve mitogenome assemblies have suggested that previously published mitogenomes were incomplete due to difficulties in assembling uncertain genomic regions (Sharbrough, et al. 2023). For instance, a partial mitogenome of *Dreissena polymorpha* (Bivalvia: Dreissenidae) was previously published, but contained a gap which targeted PCR and short-read sequencing were unable to resolve (Appeltans, et al. 2012). Recently, a complete *Dreissena polymorpha* mitogenome (67 kb) was assembled using PacBio and Oxford Nanopore sequencing by identifying the gap as a large highly repetitive segment of nearly 50k bp (McCartney, et al. 2022). Long reads also revealed that its closely related species, *Dreissena rostriformis*, has a large mitogenome (46 kb) with two blocks of repetitive DNA (Calcino, et al. 2020). In addition, similar assembly problems also have been identified and resolved in other animal lineages (Gan, et al. 2019; Formenti, et al. 2021; Kinkar, et al. 2021), accentuating the need to scrutinize published mitogenomes of bivalve using long-read sequencing and efficient workflows.

To better understand evolution of bivalves, it is essential to comprehensively explore their genetic foundation, including mitogenomes and their content. Here, we used a workflow with available long-read sequencing data to reassemble mitogenomes of mollusks, focusing on the class Bivalvia. We rechecked the assembled mitogenome structure and content, revealing unexpected and marked variability in length, number of repeats in mitochondrial URs. Moreover, we also systematically assessed the influence of different biological factors on the variability of mitogenome size. Our study suggested that long read sequencing have broad applicability to elucidating complex structure in molluscan mitogenomes and provide insights into evolution of mitogenome size in metazoans.

## Results

### Long-read Sequencing Revealed Astonishing Mitogenome Size Variation

By using a pipeline that combines long-read (PacBio and Nanopore) and short-read data sets (Fig.1A), we successfully assembled and characterized 132 complete Mollusca mitogenomes to explore mitogenome size variation across Mollusca. Additionally, 58 mitogenomes assembled based on long-read sequencing were directly obtained from the NCBI database. In total, our data contains representatives of 178 species (190 mitogenomes) belonging to 74 families, including 17 species that were not previously available in the database (Fig.1C and Table S1). Mitogenome size varied extensively across our dataset, with size varying by 7-fold, the highest being 96 kb in *Scapharca kagoshimensis* (ark shell) and the lowest being 13 kb in *Anisus vortex* (freshwater snail). Nearly three-quarters (73%) of the species had an assembled size of 13–20 kb, aligning with the typical size observed in most bilaterian animals. Of the remaining species, five-sixths have mitogenomes larger than 20 kb, all of which belong to the class Bivalvia. Comparing our assemblies with reference sequences in GenBank, we noted that many mitogenomes labeled as complete and circular were shorter than ours (Fig.1B). For instance, in the Olympia oyster (*Ostrea lurida*), our assembly revealed a 5,639-bp-long repetitive duplication, including three full-length copies of the *Cox1* gene and two copies of trnE. Similarly, reference sequences marked as partial typically result from the incomplete assembly and lack most of the URs where repeats and gene duplications typically occur. Upon ascertaining that our assemblies are likely more complete than the respective existing references, we compared size and distribution with the RefSeq representative mitogenomes in bivalves and gastropods (Fig.1D, E and Fig.2).

**Fig. 1.**
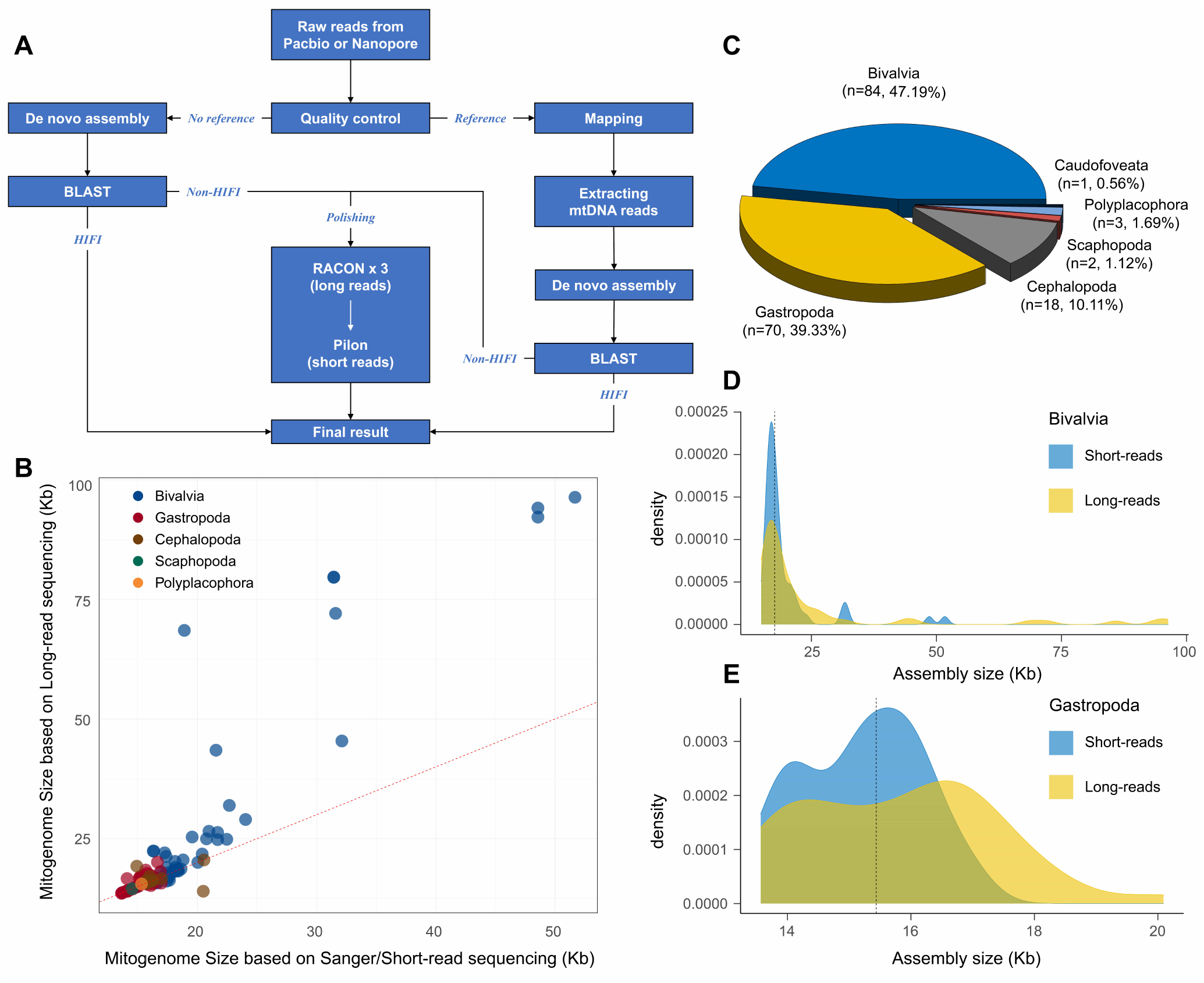
(A) Computational pipeline implemented for the mitogenome assembly based on long-read sequencing data. (B) Scatter plot of mitogenome size based on long-read and Sanger/short-read sequencing data. (C) Pie chart of mollusk data used in this study. (D) Distribution of bivalve mitogenome sequence lengths based on long-read and short-read sequencing. Length density distribution (yellow area) based on long-read sequencing and short-read density distribution (blue area).The means is the dashed line. (E) Distribution of gastropoda mitogenome sequence lengths based on long-read and short-read sequencing.

**Fig. 2.**
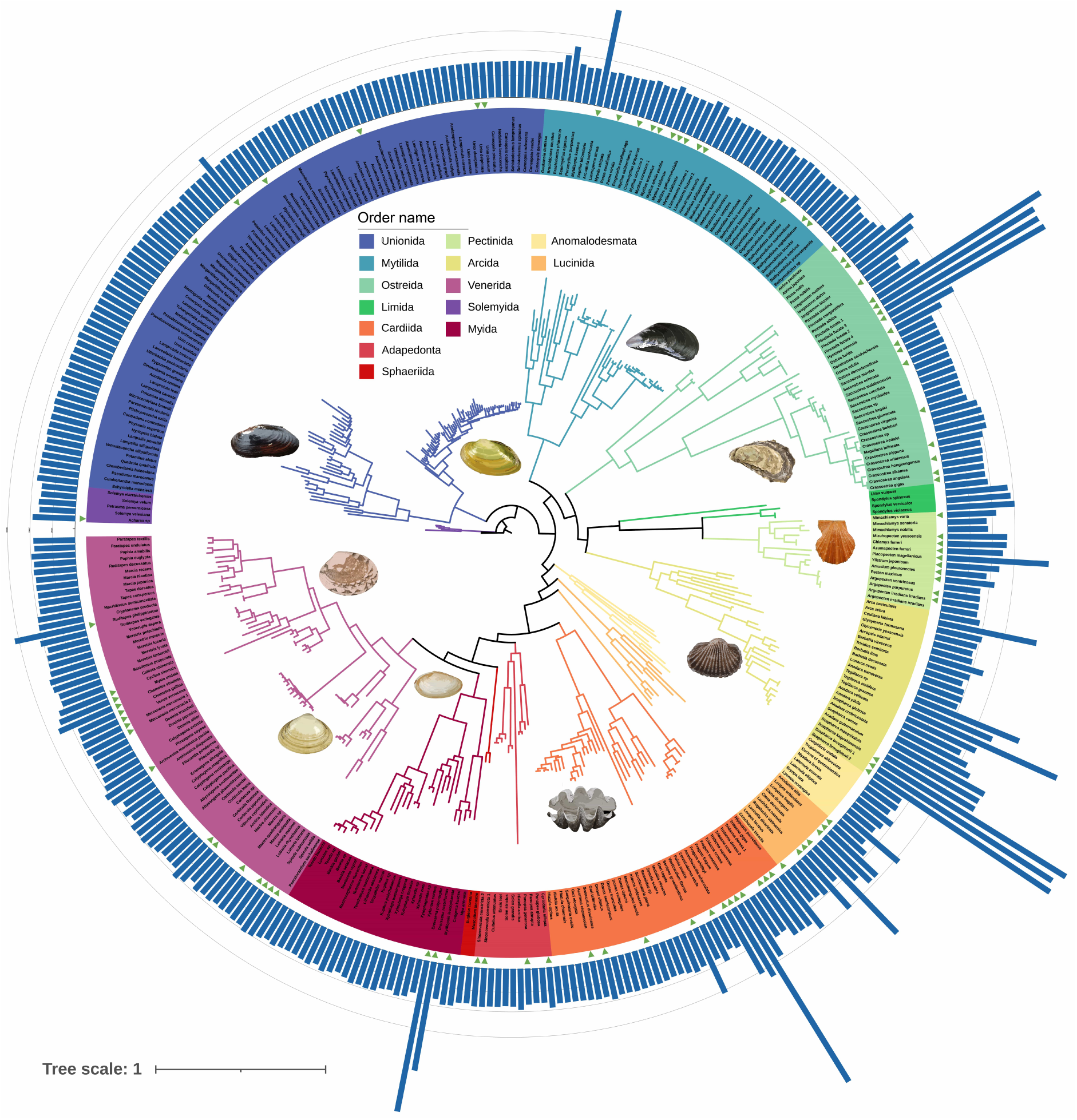
Genome-wide phylogeny of 377 bivalve species and mitogenome size distribution. Circle 1: the taxonomic names of 377 species. The color range represents the orders in Bivalvia. Circle 2: triangle represent the mitogenome assembled by long-read data. Circles 3: bule bar represent the mitogenome size of bivalve species.

For Bivalvia, 95 mitogenomes were analyzed, representing 11 orders (Table S1 and S2). We found that bivalves exhibited significant size variation compared to the mitogenomes assembled using Sanger or NGS sequencing data (Fig.S1, Wilcoxon signed-rank test, *P* < 0.001). Within these, 18 species assembled using long-read sequencing showed variation rates exceeding 20%, with the most significant increase being 262% in *Dreissena polymorpha* (freshwater mussel). Furthermore, 17 species exhibited mitogenomes surpassing 30 kb in length; among these, 7 species exceeded 50 kb, including *Pinctada imbricata* (79,602 bp) and *Scapharca broughtonii* (92,189 bp, PacBio). The ark shell (*Scapharca kagoshimensis*) stands out as the largest, with a mitogenome reaching 96,329 bp in length. Our results revealed that the existence of many complex mitogenomes that are difficult to detect with short-read sequencing. By integrating our survey of mitogenome size and the phylogeny of Bivalvia (Fig.2), we found multiple probable lineage-specific expansion events, including many not previously reported. Mitogenome expansion has independently occurred across more than 10 clades including the pearl oysters, freshwater mussels, heart cockles, giant clams, ark shells, lucinid clams, spiny oysters, and sea scallops, demonstrating that such expansion is widespread among bivalves. Particularly, the genus *Anadara*, *Pinctada* and *Fragum* exhibit the most dramatic variations in mitogenome size. In contrast, the order Unionida, despite having the largest dataset, shows minimal overall variation.

In comparison to bivalves, we observed a modest increase in the overall size of gastropod mitogenomes (Fig.1E), a difference that was statistically significant (Wilcoxon signed-rank test, P < 0.05). Notably, some gastropods exhibit considerable length variations compared to those based on first- and second-generation sequencing. For instance, the mitogenome of red abalone (*Haliotis rufescens*), as assembled by long-read data (20,092 bp), was 20.7% larger than its assembly from short-read sequencing (16,646 bp). The mitogenome of *Lottia scabra* that we assembled is 46,213 bp in length, representing the largest mitogenome identified within gastropods to date. This finding suggests a possible lineage-specific expansion within *Lottia*, as closely related species also exhibit unusually large mitogenomes (*Lottia digitalis*, 26,835 bp). Despite the occurrence of large mitogenomes in some gastropod groups, both their maximum size and prevalence are considerably lower compared to those observed in bivalves. Beyond two classes, most mollusk species across other classes share the typical bilaterian mitogenome sizes, ranging from 15 to 20 kb (Table S1 and Fig.S2). While a few cephalopod species may exceed 20 kb, the increase is not substantial. Additional data collection is necessary to further investigate these groups.

### Factors that Influence Bivalve Mitogenome Size

To investigate the causes of size variation, we conducted analyses at various levels, considering factors external to mitogenome, such as environment, lifestyle, and nuclear genome, as well as internal factors like mitochondrial gene duplication, GC content, repeated sequences, and potential functional elements.

### Nuclear Gnome Size, Ecological Factors and Lifestyle

Mitogenome vary conspicuously in size across Mollusca. Similarly, there was substantial variation in nuclear genome size, which varied by about 15-fold across our dataset, from 0.4 Gb in *Chrysomallon squamiferum* (scaly-foot snail) to 6 Gb in *Pictodentalium vernedei* (scaphopod). A potential correlation between mitochondrial and nuclear genome sizes might exist due to mito-nuclear interactions and NUMT insertions (Puertas and González-Sánchez 2020; Weaver, et al. 2022). However, our results revealed that, in both bivalves and gastropods, there is no significant correlation between the sizes of mitochondrial and nuclear genomes (Fig.3F, Bivalve, Kendall’s rank, tau = -0.14, P > 0.05; Fig.S3, Gastropod, Kendall’s rank, tau = 0.09, P > 0.05). This suggests that mitochondrial and nuclear genome sizes may have evolved independently. Additionally, based on our collected data (Table S3), environmental factors, distribution and lifestyle appear to exert no influence on mitogenome size. Species exhibiting mitogenome expansion (e.g. pearl oysters, ark shells, freshwater mussels, cold water scallops) inhabit diverse environments and display varied lifestyles. For example, *Tegillarca granosa* (ark shell) live in temperate intertidal zones, while *Pinctada fucata* (pearl oyster) is prefers warm waters and attaches to rocks or other hard substrates. *Fragum sueziense* (heart cockle), which forms photosymbiosis, can be found up to 20 meters deep, buried in sandy seabed. Strikingly different, *Ctena decussata*, known for chemosymbiosis, is benthic with a depth range of 125-200 meters. These examples suggest no consistent environmental or lifestyle trait linked to variations in mitogenome size, though further work is needed to validate this observation.

**Fig. 3.**
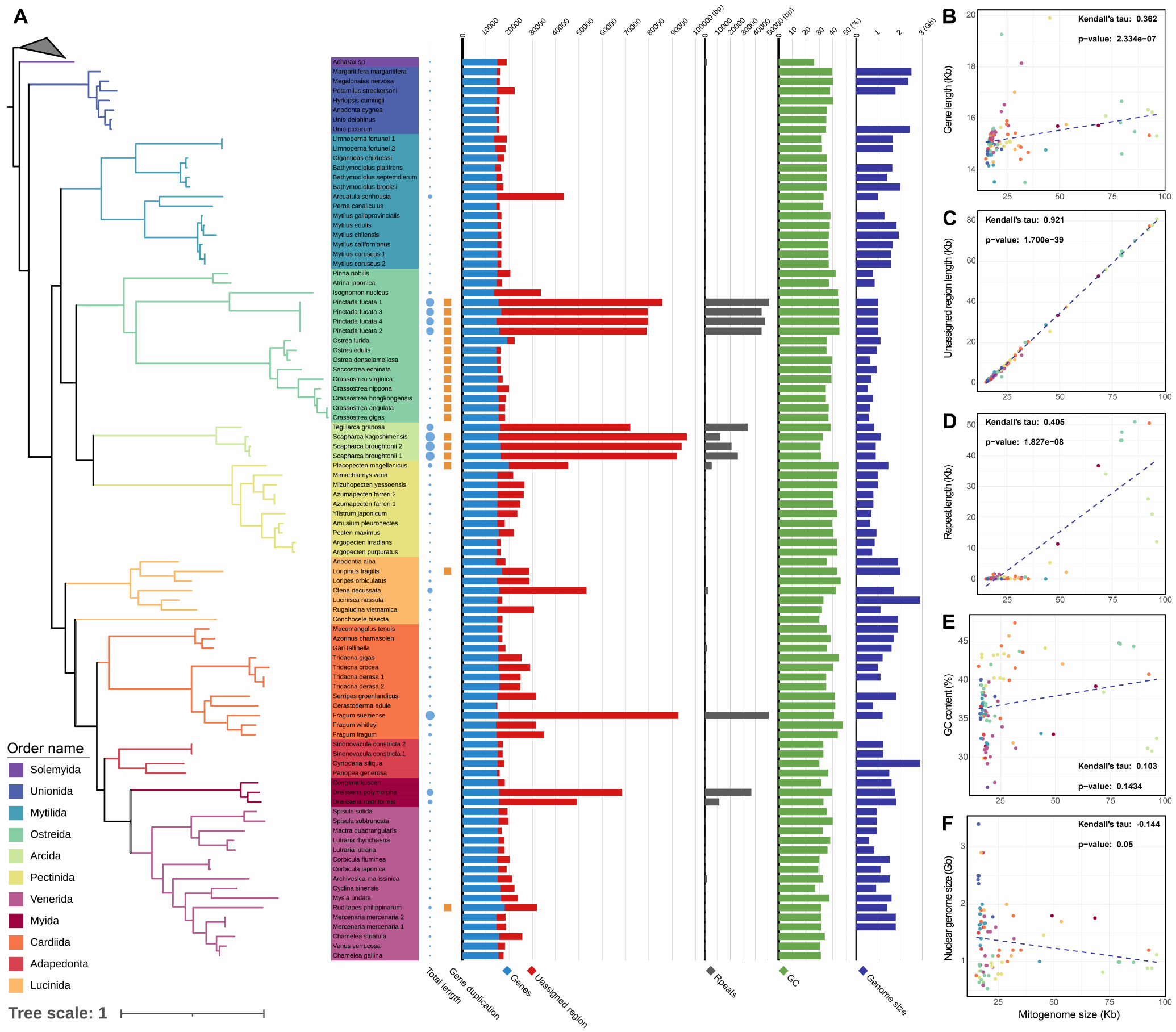
Phylogenetic and mitogenomic comparisons among 94 bivalve species. (A) From left to right: the phylogeny built from 12 mitochondrial genes using Maximum Likelihood Inference. The branch color represents the orders in Bivalvia; mitogenome size and gene duplication; coding region (blue) and unassigned region (red); repeats; GC content; Genome size. Data for the bar plot can be found in Supplementary Table S4. (B-F) The correlation of the factors (Genes, URs, repeats, GC content and nuclear genome size) on mitogenome size variation across Bivalvia, respectively.

### Gene Duplication

Gene duplication is a crucial evolutionary mechanism that plays a vital role in the evolution of mitogenome complexity (Formenti, et al. 2021; Ghiselli, Gomes-dos-Santos, et al. 2021). In bivalves, variations in gene length are primarily attributed to gene duplications and changes in the length of protein coding genes. Our analysis demonstrates that the length of genes does not strongly correlate with the length of mitogenomes (Fig.3B, Kendall’s rank, tau = 0.36, P < 0.05), suggesting that variations in mitochondrial gene length do not account for all observed expansions. Although significant expansions in two groups (*Scapharca* and *Pinctada*) were accompanied by gene duplication, this does not imply a causal relationship, as gene duplication is not the primary cause of these expansions (Fig.3A). Conversely, for clades without significant mitogenome expansion, such as in Ostreidae (e.g. *Ostrea lurida*), gene duplication appears to be a key factor driving expansion. Nonetheless, the contribution of gene duplication to mitogenome size cannot be ignored. In the mitogenome of *Placopecten magellanicus*, we discovered a block duplication of four genes (*Atp6*-*Cob*-*Cox2*-*Cox3*), which reached over 3000 bp. This duplication was completely absent in the reference sequences in Genbank (DQ088274), strengthening the notion that Sanger/short-read assemblies inevitably fail to represent gene and segment duplications. Moreover, a potential trend was observed where similar types of genes are duplicated within closely related taxa. Specifically, duplications of the rRNA 16S gene were identified in Ostreidae, while duplications of the rRNA 12S gene were found in the closely related clade, Pteriidae. Concerning tRNAs, species with significant mitogenome expansions often have an increased number of tRNAs (Table S4), though their overall impact on genome size is minimal.

### Repeats in Unassigned Regions

Previous analyses have shown that bivalves possess a significantly higher proportion of URs compared to other groups, exhibiting the highest median percentage of URs among Metazoans (Ghiselli, et al. 2013). Our findings corroborate this and reveal a strong, positive correlation between the length of URs and mitogenome size (Fig.3C, Kendall’s rank, tau = 0.92, P < 0.05). Taking into account all the correlation analyses performed, it is evident that URs are a major contributor to the expansion of mitogenomes. Additionally, we have found that repeats also display a similar trend; repeat sequences vary widely among most bivalves, with their lengths showing a moderate positive correlation with mitogenome size (Fig.3D, Kendall’s rank, tau = 0.4, P < 0.05). Based on the annotation results, three types of repeat sequences were identified: simple repeats, low complexity repeats, and repeats from unknown families. Overall, over 70% of assemblies (67/92) contained between 1 to 264 repeats, most of which is simple repeats. Generally, the number of repeats increased with increasing mitogenome size, although there was a tendency for the number of repeats to decrease at medium mitogenome sizes. We found that in larger mitogenomes, there is a diversity of repeat types, including many from unknown families. In contrast, smaller mitogenomes contain only simple repeats or none at all.

Moreover, the results show that repeat length is not strongly positively correlated with UR length (Kendall’s rank, tau = 0.42, P < 0.05), indicating that these repeats represent just one of multiple factors contributing to UR expansion in bivalves. In larger mitogenomes, typically those over 60 kb, many unknown repeat families are found, constituting a considerable proportion of URs. Conversely, in mitogenomes ranging from 30 kb to 60 kb, despite comparable expansions, these repeat families are absent; only simple and low complexity repeats are detected, and they occupy a very small proportion of the URs, with most of the area remaining unidentified. For example, the UR of lucinid clam *Rugalucina vietnamica* is over 15 kb in length, of which only 256 bp of simple repeats are found. Furthermore, in some species with expanded mitogenomes, such as *F. fragum* and *F. whitleyi* (both exceeding 30 kb), no repetitive sequences were observed at all, hinting at the potential presence of additional content. Species undergoing expansion without numerous repeat sequences may possess other functional components.

### GC Content

One of the important qualitative aspects of genomic architecture is the genomic nucleotide composition, commonly quantified by the proportion of guanine and cytosine bases in the DNA molecule (GC content) (Šmarda, et al. 2014). To further explore potential compositional drivers of mitogenome size, we examined the variation in GC content across species, which spanned from 26.09% in *Acharax sp.* to 47.33% in *Fragum whitleyi*. The results show no significant correlation between mitogenome size and GC content (Fig.3E, Kendall’s rank, tau = 0.1, P > 0.05), suggesting that GC content does not preferentially associate with mitogenome size. While previous studies (Sun, et al. 2015b; Sun, et al. 2015a; Sun, et al. 2015c) have characterized mitochondrial URs as AT-rich, our findings revealed no significant correlation between GC content and the length of URs (Kendall’s rank, tau = 0.11, P > 0.05). This inconsistency may be attributed to incomplete assembly. Alternatively, some variation in GC content may be driven by the density of features such as repeats (Wright, et al. 2024). Contrary to expectations, although larger mitogenomes tend to have a higher number and size of repeats, no significant correlation was observed between GC content and repeat length in these mitogenomes (Kendall’s rank, tau = - 0.03, P > 0.05).

### Potential Functional Elements in Unassigned Regions

Recent studies have shown that some mitogenomes contain multiple forms of functional elements in addition to traditional genes, which is not surprising given the importance of mitochondrial activities in multifaceted cellular functions (Zhu, et al. 2022; Kienzle, et al. 2023). To explore this further, we investigated non-coding RNAs (ncRNAs) and protein-coding genes in complete mitogenomes to predict potential functional components. Using cmsearch from Infernal to search the assembled mitogenomes against the Rfam database, we identified matches for 29 ncRNAs in 22 bivalve species (Fig.4 and Table S5). Notably, almost all of these matches are miRNAs, with the exception of two sRNAs. All of these RNAs are located in mitochondrial URs (Table S2). Among the species studied, the mitogenome of *Dreissena rostriformis* exhibited the greatest diversity of ncRNAs. Additionally, mir-5681 was the most frequently observed ncRNA across various species. Some ncRNAs, such as mir-627, were present in multiple copies within the mitogenome, indicating a pattern of content duplication in the URs. To broaden our understanding, we conducted an ncRNA survey on over 500 bivalve mitogenomes assembled by Sanger or short-read sequencing (Fig.S4). Our results identified matches for 62 ncRNAs in 70 species, with mir-3149 and mir-9555 being the most frequently occurring. However, no large-scale duplication of ncRNAs was observed. In addition, we extended our survey to include more than 800 gastropod mitogenomes available in the NCBI database (Fig.S5 and Table S6). This analysis identified matches for 64 ncRNAs in 126 species. Among these, mir-297 was the most prevalent, detected in 35 species. For protein detection, we analyzed over 500 bivalve mitogenomes, including those assembled using third-generation, Sanger, and short-read sequencing techniques. Matches for 29 proteins were identified across 20 species (Fig.S6). These sequences were all located in URs and exhibited varying degrees of duplication. Although our findings do not conclusively prove the existence of these functional elements, they suggest that URs in bivalve and gastropod mitogenomes may harbor a significant number of functional elements. This potential contributes to the complexity of mitogenome architecture.

**Fig. 4.**
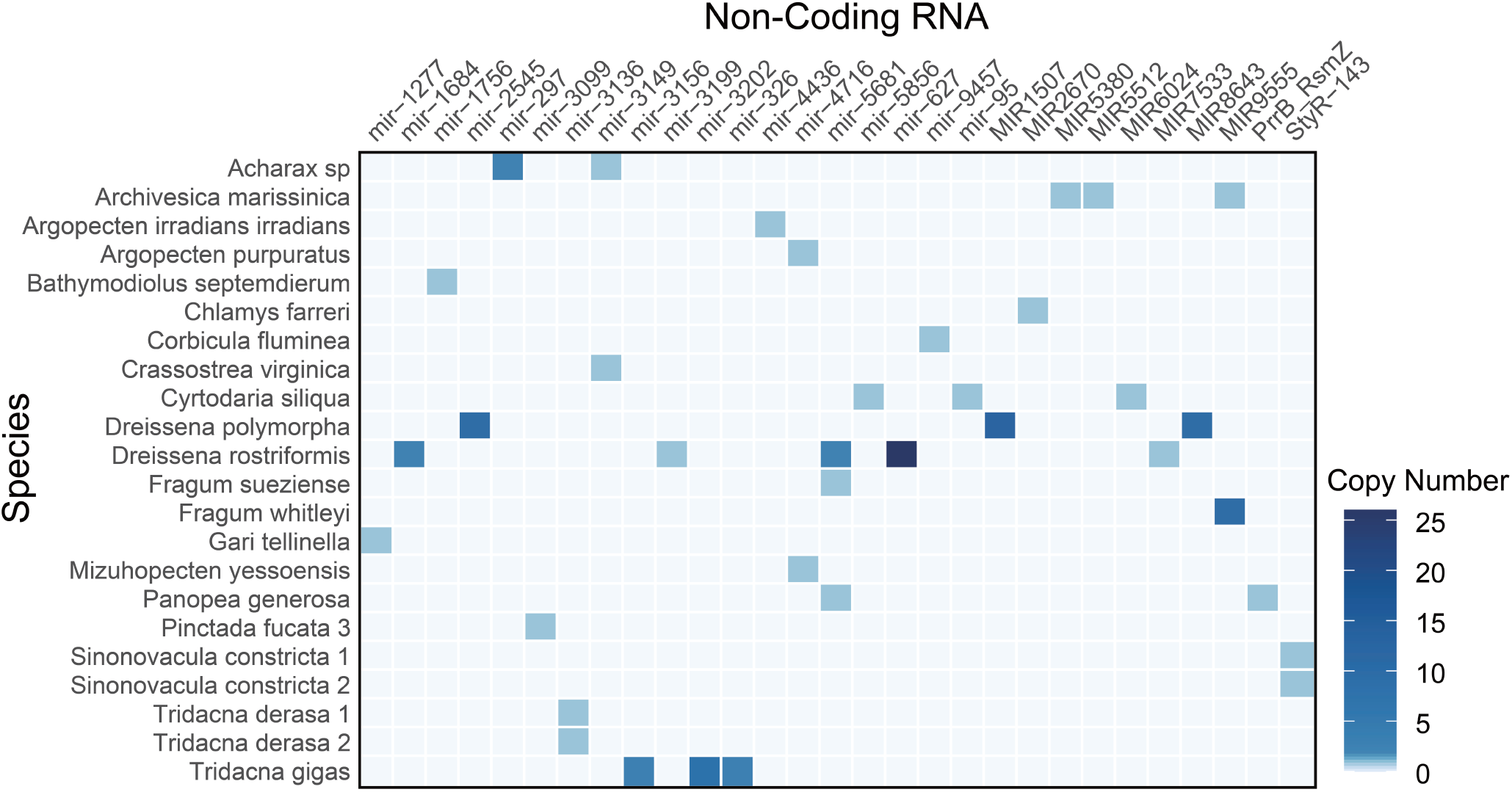
Non-coding RNA detected in the complete mitogenomes of bivalves. The color depth represents the change in copy number.

### Distribution and Rearrangements of Expansion Regions across Bivalves

Complete mitogenomes were determined for 85 bivalve species included in this study. Our annotations allowed us to establish the order of protein-coding genes, rRNA genes, and expansion regions. To examine how the mitochondrial genome expands, we analyzed the locations of expansion regions to identify potential preferences for expansion sites. However, extensive rearrangements within bivalve mitogenomes are so pronounced that identifying consistent patterns or preferences across different lineages remains challenging (Boore 1999; Sun, et al. 2020; Malkócs, et al. 2022). Consequently, we focused on closely related species exhibiting only minor rearrangements to further investigate patterns of expansion.

The comparative analysis of mitochondrial synteny across multiple lineages has uncovered that expansion regions do not exhibit a preference for specific genes (Fig.5). The number of expansion regions identified ranged from one to three. Currently, the observed patterns among closely related species can be classified into two distinct types. First, species display an identical number of expansion regions located at the same positions. In these cases, the size differences within or between species are primarily attributable to variations in the lengths of these expansion regions. For instance, in the mitogenomes of three *Tridacna* species, the synteny of genes and expansion regions remains conserved, with an UR consistently occupying the same position. Variations in the lengths of these URs contribute to the observed differences in mitogenome lengths, suggesting that the variation of extended URs may not influence the rate of gene rearrangement. Second, although some closely related species do not maintain a consistent number of expansion regions, at least one expansion region is invariably found in a common position. For example, two *Dreissena* species share a UR at the same locus, yet the total number of URs varies. Specifically, *D. rostriformis* possesses an additional UR of 5,705 bp in its mitogenome. Similarly, within the family Arcidae, three species demonstrate high gene and UR collinearity, with a novel UR appearing in *T. granosa*. This indicates that the emergence of URs may not impact gene rearrangement. Conversely, gene rearrangement may influence the generation of URs. In *D. rostriformis*, the position of the *Nad4L*-*Cox3* gene cluster shifted to the position of UR2 relative to *D. polymorpha*, and a new UR emerged at the original *Nad4L*-*cox3* location. This suggests that sites of gene rearrangement may be more susceptible to the formation of expansion regions, warranting further investigation. Additionally, we observed a clear demarcation between coding regions and URs. Coding genes are typically clustered into one or two groups and remain together even as the mitochondrial genome expands.

**Fig. 5.**
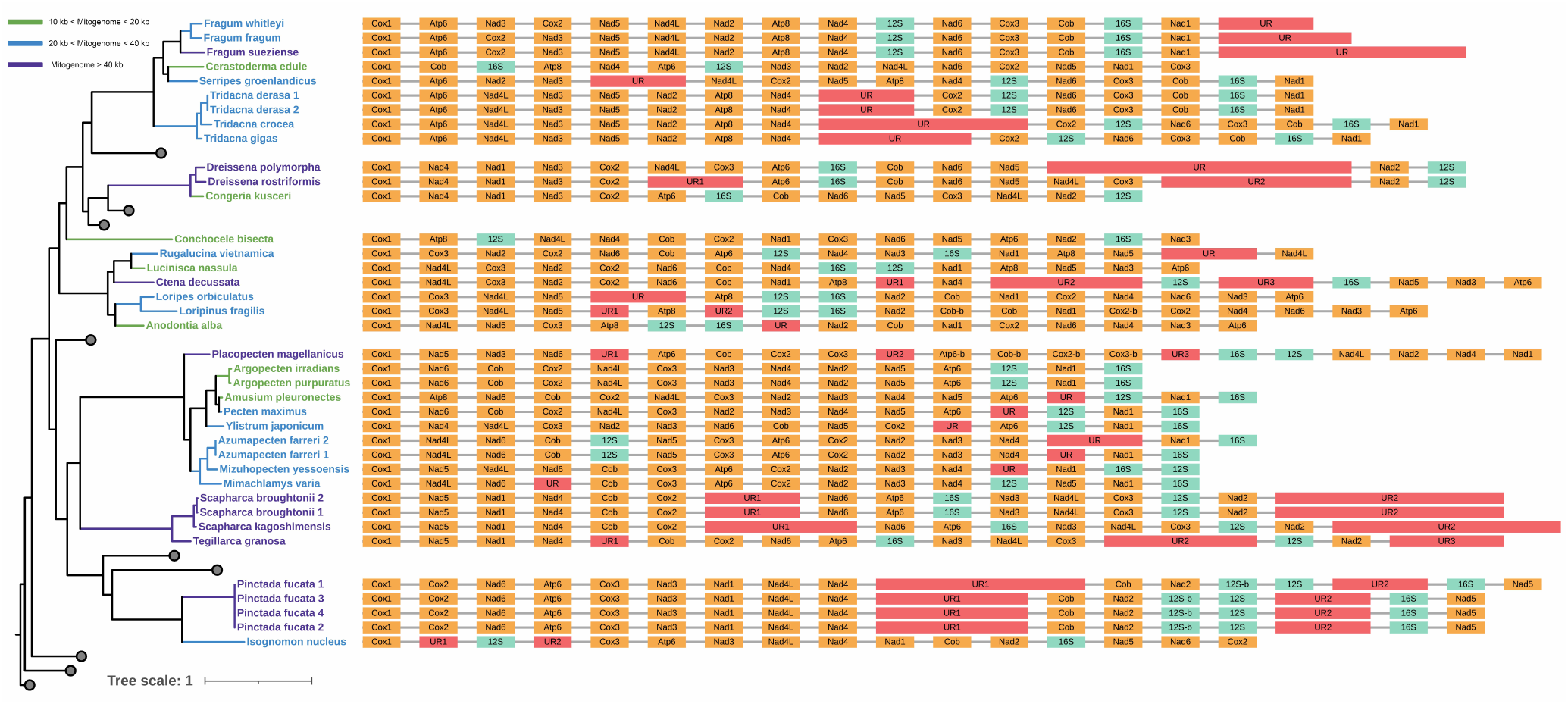
Phylogenetic relationships of the bivalve species with large mitogenomes and the distribution of large-scale expansion events.

### Bivalvia Mitogenome Phylogeny and its Limitation

To examine the distribution of mitogenome sizes, we constructed a phylogenetic tree of 377 bivalve species based on 12 mitochondrial genes, covering most major orders (Fig.2). Although this tree is currently the most comprehensive based on mitochondrial genomic dataset, the backbone of the bivalve tree of life has recently been extensively studied by transcriptomic and genomic data (Lemer, et al. 2019; Song, et al. 2023). Therefore, we briefly compare the differences with previous studies, highlighting the limitations of mitochondrial genes construction. Our phylogenetic analysis supports the monophyly of most high-level clades including Protobranchia, Palaeoheterodonta, Pteriomorphia and Euheterodonta. Notably, Palaeoheterodonta appeared as sister group to Pteriomorphia and Heteroconchia, which is inconsistent with the transcriptomic and genomic trees. For Pteriomorphia, Mytilida did not cluster with Ostreida (Pinnidae, Ostreoidea and Pterioidea) but instead appeared as sister group to all other pteriomorphians. Limida was not found to be a monophyletic group. For Euheterodonta, our analyses recovered a similar topology to that of the transcriptomic tree. However, Lucinida was found to be a polyphyletic group. Additionally, we found that selecting outgroups exclusively from either Gastropoda or Scaphopoda greatly influences the inferred phylogenetic relationships within Pteriomorphia (Fig.S7). These observations indicate that phylogenetic tree construction by mitochondrial genes may have inherent issues and limitations specific to bivalves, potentially hindering exploration of deep evolutionary relationships.

## Discussion

### Long-Read Sequencing in Unraveling Complexity of Mitogenomes

Long-read sequencing has greatly improved the quality of mitogenome assembly. The majority of published large mitogenomes are linear and incomplete, suggesting challenge of assembling complex mitogenomes. Compared to existing reference mitogenome assemblies, our assemblies filled gaps larger than 1000 bp in 36% of cases, added missing repeats, and identified additional functional elements or gene duplications, all of which led to novel discoveries using these complete assemblies (Table S1, S2 and S4). For instance, a very recent publication employing only short-read sequencing (Illumina) reported a complete, circular mitogenome for the scallop *Ylistrum japonicum* with a total length of 19475 bp (PP571649). In contrast, our assembly of *Y. japonicum*, utilizing long-read sequencing data, unveiled a mitogenome size of 23,580 bp, more than 4 kb larger than the assembly produced without long-read sequencing technology. Our results present numerous instances of such discrepancies, suggesting that despite the extensive publication and deposition of mitogenome sequences in public databases, there is significant evidence of incompleteness, primarily due to limitations of traditional sequencing techniques in accurately capturing repetitive regions.

Bivalves can be regarded as a model system for studies focusing on evolution of mitogenome structure and content. We systematically examined size and structure in mitogenomes of diverse bivalve and found that multiple instances of extreme size and lineage-specific expansion, suggesting the repeated evolution of clades with large and small mitogenomes across bivalve diversification (Fig.2). The observed variability and frequent occurrences point to instability and considerable plasticity within the UR of some bivalve mitogenomes. In particular, our reported the mitogenome of the blood clam *Scapharca kagoshimensis* (96 kb) is one of the largest among bilaterian animals published to date, highlighting a more dramatic UR expansion within Arcidae than previously documented (Zhang, et al. 2022). Based on available data, bivalves exhibit more frequent expansion and significant mitochondrial length variation compared to other mollusks and even other bilaterians (Fig.2 and Fig.S2). Long-read sequencing can uncover many previously undetected mitogenome variations, particularly in unique groups like bivalves, where additional variations are likely to be discovered in the future.

The mitogenome is often overlooked in large genome sequencing projects, despite the importance of mitochondrial variation for medicine and studies of animal molecular systematics and evolution (Morgan, et al. 2022; Meadows, et al. 2023). During our data collection, we noted that many sequencing projects primarily targeted nuclear genomes, neglecting assembly and publication of mitogenomes using long-read sequencing. So far, only the Darwin Tree of Life Project has published mitogenomes assembled by long-read data, specifically limited to the PacBio HIFI data within the project (Consortium 2022). With advances in sequencing technology and explosion of data, we call for increased focus on complete assembly of mitogenomes using third-generation sequencing. Moreover, through comparisons among Nanopore data, PacBio continuous long-read (CLR) data, and PacBio high-fidelity (HiFi) data, it was established that PacBio HiFi data boasts remarkable accuracy, ensuring high-quality assembly. This is evidenced by the detailed annotation of the mitogenomes, which does not need further polishing with short-read data. Long-read sequencing data, especially PacBio HiFi data, should serve as the foundational resource for mitogenome research to prevent incomplete assemblies, thus enabling more comprehensive studies on mitogenomes.

### Evolutionary Patterns of Bivalve Mitogenome Expansion

The evolution of mitogenome size in bivalves appears to be primarily driven by molecular factors. In our study, we investigated several external factors, including nuclear genome size, environmental conditions, and lifestyle, that could potentially influence the size of bivalve mitogenomes (Struck, et al. 2023). However, our results revealed no evidence to suggest that mitogenomes adapt to these types of selective pressures. At the molecular level, the significant variation in UR sequences across different bivalve species indicates that these molecular features play a crucial role in determining mitogenome size. This finding suggests that the evolution of mitogenome size is more likely influenced by intrinsic molecular evolutionary processes rather than by life history traits or ecological factors.

The length of the unassigned region (UR) is the primary factor determining mitochondrial genome size, and rapid evolution may account for the observed variation in UR length. For example, in the genera *Tridacna*, *Fragum*, and *Scapharca*, the length of the UR varies greatly among closely related species (Zhang, et al. 2022), while other parts of the mitogenome show little variation. Even within the same species (e.g., *S.broughtonii*, *P. fucata* and *Azumapecten farreri*), the length of the UR can vary considerably, indicating that the structure and content of URs evolve rapidly. We identified a variety of contents within URs, including repetitive sequences, potential RNA or protein structures. Regardless of whether the sequences are repetitive or represent other functional elements, we observed a prevalent trend of duplication within the UR contents of most species. This suggests that duplication may be the primary pattern of mitochondrial UR expansion and evolution in bivalves.

The expansion of the mitogenome in bivalves has provided the capacity for the generation of functional elements. According to previous studies (Lee, et al. 2013; Breton, et al. 2014; Lu, et al. 2024), besides traditional genes, the mitogenome appears to be under evolutionary pressure to expand its functions and incorporate additional functional elements. In the human mitogenome, although its length remains highly conserved and similar to that of most bilaterians, at least nine additional protein genes have been identified (Kienzle, et al. 2023). These functional sequences frequently overlap, sometimes completely, with other coding genes, leading to the possibility that a single sequence mutation could impact multiple genes (Breton 2021). In certain bivalve species, proteins found within the mitochondrial URs may play a role in sex determination (Breton, et al. 2011; Milani, et al. 2013; Ouimet, et al. 2020). Given the abundance of meaningful components in URs, we speculate that the expansion of the mitogenome may provide the necessary space for the generation of functional elements. However, the roles and significance of the contents within these URs remain to be thoroughly investigated.

### What Determines the Size of Bivalve Mitogenomes?

The extreme size variation in bivalve mitogenomes may result from the relaxation of strong selective pressures, such as those associated with innate immunity. Traditionally, it has been believed that the selection pressure on mtDNA is primarily driven by its role in energy metabolism. However, recent studies have highlighted that mtDNA also plays a pivotal role in activating innate immune responses and inflammation (Riley and Tait 2020; Kim, et al. 2023; Newman and Shadel 2023). These functions of mtDNA suggests that evolutionary pressures on mitogenome are more complex than previously thought. The mechanisms involved in mtDNA release, such as the formation of mitochondrial pores, and the interaction with DNA sensors of the innate immune system (e.g., the cyclic GMP-AMP synthase), may influence the size and structure of the mitogenome. Furthermore, mt-Z-DNA (Z-form mtDNA) released from mitochondria is stabilized by Z-DNA binding protein 1 (ZBP1), an innate immune receptor, to sustain interferon signaling, suggesting that the helicity of mtDNA structure may also subject to selection (Lei, et al. 2023). These interactions could serve as selective forces that constrain the evolution of the mitogenome, ensuring it remains efficient for immune functions. Interestingly, bivalves exhibit unique immune characteristics. For instance, they present the largest and most diversified repertoire of Toll-like receptors in the animal kingdom, which function as pattern recognition receptors in the innate immune system (Saco, et al. 2023). Some Toll-like receptors (e.g. TLR9) can engage mtDNA that is released into the cytoplasm or extracellular space under conditions of cellular stress and mitochondrial dysfunction (Marchi, et al. 2023). Moreover, some bivalves have the ability to synthesize, store, and secrete the antibiotic erythromycin to combat bacterial infections (Yue, et al. 2022). Several marine species have been found to host eight transmissible cancer lineages, which probably spread via the transfer of free-floating cells in seawater (Bruzos, et al. 2023). These immune peculiarities suggest that bivalves have adapted their immune systems in ways that are distinct from other animals. Such abnormalities and the corresponding variations in the mitogenome of bivalves may be interconnected. Therefore, we hypothesize that the relaxation of selective pressure on mitogenome size in bivalves may be attributed to changes in the innate immune pathways related to mtDNA. This provides a new perspective on why many bivalves can maintain such large mitogenome sizes. Further research into these mechanisms could offer valuable insights into the evolutionary pressures on mitogenomes in bilaterians.

## Methods

### Data Collection and Genome Sequencing

We collected all long-read sequencing data available for mollusks from the SRA database, as well as short-read sequencing data that can be used for mitogenome polishing. This data includes 84 species (95 mitogenomes) of bivalves, 70 species (71 mitogenomes) of gastropods, 18 species of cephalopods, 3 species of polyplacophorans, 2 species of scaphopods and 1 species of caudofoveata (Table S1). In addition, we sequenced a new individual of blood calm *S. broughtonii* and generated 32.5 Gb of PacBio long-read data. For Illumina sequencing, a paired-end (PE) library with an insert size of 450 bp was constructed and sequenced with an Illumina HiSeq X Ten platform. A total of 96 Gb Illumina data were generated and used for mitogenome polishing. High-quality DNA was used for library preparation and high-throughput sequencing using PacBio and Illumina platforms (Tianjin Novogene Bioinformatics Technology Co. Ltd., Beijing, China).

### Mitogenome Assembly and Annotation

To obtain complete mitogenomes of Mollusca species, PacBio and Nanopore long-reads from SRA database were aligned to the mitogenome of the species or its closely related species using Minimap2 v2.17 (Li 2018). The aligned reads were then extracted using Samtools (Li, et al. 2009) and subjected to error correction, trimming, and assembly using Canu v1.7 (Koren, et al. 2017). In cases where the mitogenome of the species or its closely related species had not been published, the long-reads were directly de novo assembled without mapping, using an approximate genome size of 1 Gb. Contigs meeting the criteria “suggestRepeat=no, suggestBubble=no, suggestCircular=yes” were selected. If the resulting draft assembly contained more than one contig, a blast search was conducted to confirm which contig contained all mitochondrial genes. Due to the circular nature of the mitogenome, the contig often consists of long concatemers of mtDNA sequences, which we manually annotated as a single circular mitogenome. The putative mitochondrial contig was polished using three rounds of Racon v1.4.13 (Vaser, et al. 2017) with long-read data, followed by a round of polishing using short-read data. The short reads were mapped with BWA (Li and Durbin 2010) and the alignment file was piped into Pilon v1.22 (Walker, et al. 2014) to specifically correct for small indels, gaps and SNPs. The depth of coverage throughout each mitogenome was calculated with Bedtools (Quinlan and Hall 2010) and Bamdst (https://github.com/shiquan/bamdst). We did not perform the aforementioned polishing steps on contigs assembled by HIFI data, because HiFi sequencing generates extremely accurate long-reads. The complete procedure for the mitogenome reconstruction is schematized in Figure 1A.

The polished mitogenomes and the published mitogenomes without annotation were initially annotated with the MITOS2 webserver using the invertebrate genetic code. The boundaries of protein-coding genes were further annotated using the ORF Finder (https://www.ncbi.nlm.nih.gov/orffinder/) and manually inspected with the published mitogenome as the reference. Then, ribosomal RNA (rRNA) genes were identified using BLAST (Altschul, et al. 1997) and their boundaries were determined by identifying the start and stop sites of adjacent genes. Each final circular assembly was rotated to begin at the *Cox1* gene. The graphical assembly of contig flagged as circular by Canu was visualized on Bandage v0.8.1 (Wick, et al. 2015). Annotation information for the mitogenomes is in Supplementary Table S2.

### Characterization of Repeats

Repeat sequences of the assembled mitogenomes were identified through a combination of de novo and homology-based methods. Species-specific consensus sequence libraries were generated using RepeatModeler v2.0.2 (Flynn, et al. 2020), which integrated the outputs of two distinct algorithms: RepeatScout v1.0.6 and RECON v1.08 for de novo detection. Subsequently, RepeatMasker v4.1.2 (https://repeatmasker.org) was utilized to screen and classify these repetitive sequences against our de novo consensus sequence library and the Dfam database by the comparison program of RMblast. The resulted unknown repeat families were combined with the default full RepeatMasker database.

### Investigation of Potential Functional Elements

To search for functional elements in the mitogenomes beyond traditional genes, we conducted a preliminary search for non-coding RNAs (ncRNAs) and protein-coding genes. The complete mitogenome sequences were used for ncRNA search and annotation using Infernal v1.1.3 cmsearch (Nawrocki and Eddy 2013) against the Rfam database (Kalvari, et al. 2021). Results with an E-value of less than 1 × 10^-6^ were retained (Fremin and Bhatt 2021). All tRNA and rRNA sequences were excluded from the analysis, and the lowest-scoring overlaps identified by cmsearch were removed. For the identification of protein-coding genes, we utilized all 6-frame translations of the mitogenomes generated by getorf (with parameters -find 1 -minsize 30) from the EMBOSS v6.6.0 (Rice, et al. 2000). These translations were subsequently screened against the Pfam database using Hmmscan v3.3 (Finn, et al. 2011) to predict protein homologous sequences. Results with an E-value less than 1 × 10^-5^ were retained. All results pertaining to conventionally encoded mitochondrial genes were removed. The visualization of non-coding RNAs and protein-coding genes was performed in R using the ggplot2 and reshape2 packages.

### Determine External and Intrinsic Factors Influencing Bivalve Mitogenome size

For intrinsic factors, mitochondrial annotation data were used to quantify the lengths and counts of coding genes, tRNAs, and rRNAs, along with their aggregated lengths and totals. The overall size of URs was estimated by subtracting the total gene length from the mitogenome size. Gene duplications were identified through BLAST results and gene occurrence tallies, and GC content was analyzed using EMBOSS tools. Repeat lengths were assessed based on repeat characterization data. Regarding external factors, genome size information for each species was gathered from public databases. Data on habitat and lifestyle were collected from the literature and further completed based on the authors’ observations. These factors will be systematically analyzed to elucidate their potential influence on the variability of mitogenome sizes across bivalve species.

### Statistical Significance and Correlation Analyses

To assess the differences between long-read and Sanger/short-read sequencing assemblies, we gathered existing Sanger/short-read sequencing references for the species under study. For species with multiple mitogenomes available, only the largest assembly was retained for analysis. The Wilcoxon test was applied in R to determine the significance of assembly size differences, focusing primarily on Bivalvia and Gastropoda due to the scarcity of samples from other classes. To identify which mollusks with significant mitogenome size variations after long-read sequencing, we established a stringent threshold—mean plus two standard deviations. Species surpassing this threshold were classified as having significant changes in mitochondrial genome size. Due to the data set’s deviation from a normal distribution, we utilized outlier analysis to identify species with notably larger mitochondrial genomes, considering Z-scores above 2 as significant. To understand the relationship between mitogenome size and various intrinsic and external factors, comprehensive correlation analyses were conducted in R. We conducted correlation tests to determine the strength and significance of relationships between these factors and mitogenome size, employing the Kendall’s tau coefficient to quantify linear associations.

### Generate Limited Reference Tree for Comprehensive Analysis of Mitogenome Size Variations

Despite the limitations of building phylogenetic trees with mitochondrial genes, to show the relationship between mitogenome size and lineage, we used 377 mitogenome sequences (94 mitogenomes assembled by long-read data and 283 mitogenomes downloaded from Genbank) covering 14 orders to reconstruct the phylogenetic tree of Bivalvia (Fig.2). Three species from Scaphopoda (*Siphonodentalium dalli*, *Pictodentalium vernedei*, *Graptacme eborea*) were used as outgroups based on data availability and Mollusca evolutionary history (Song, et al. 2023). The amino acid sequences of 12 protein coding genes (excluding atp8) were individually aligned using Mafft 7.475 (Katoh and Standley 2013) and then poorly aligned positions were removed in TrimAl 1.4 (Capella-Gutiérrez, et al. 2009) under the “automated1” setting. Gene alignments were subsequently concatenated into a matrix using FASconCAT (Kück and Longo 2014) and analyzed under maximum likelihood using IQ-TREE v2.0.3 (Nguyen, et al. 2015). An optimal substitution model was automatically selected, whose robustness was assessed with 1000 replicates of ultrafast bootstrapping. Additionally, we construct the phylogenetic tree of bivalves assembled by long-read sequencing to present information of their mitogenome features. We also used different outgroups to build maximum likelihood trees to discuss the impact on the phylogeny of Bivalvia. All generated trees were displayed in Figtree and annotated using iTOL (Letunic and Bork 2021).

## Acknowledgments

We thank the support of the High-Performance Biological Supercomputing Center at the Ocean University of China for this research. We also acknowledge Yi Liu for assistance with guidance and refinement of figure preparation.

## Data availability

All high-throughput sequencing data have been deposited at the NCBI SRA database under BioProject PRJNA1191059.

## Supplementary files

**Fig.S1.**
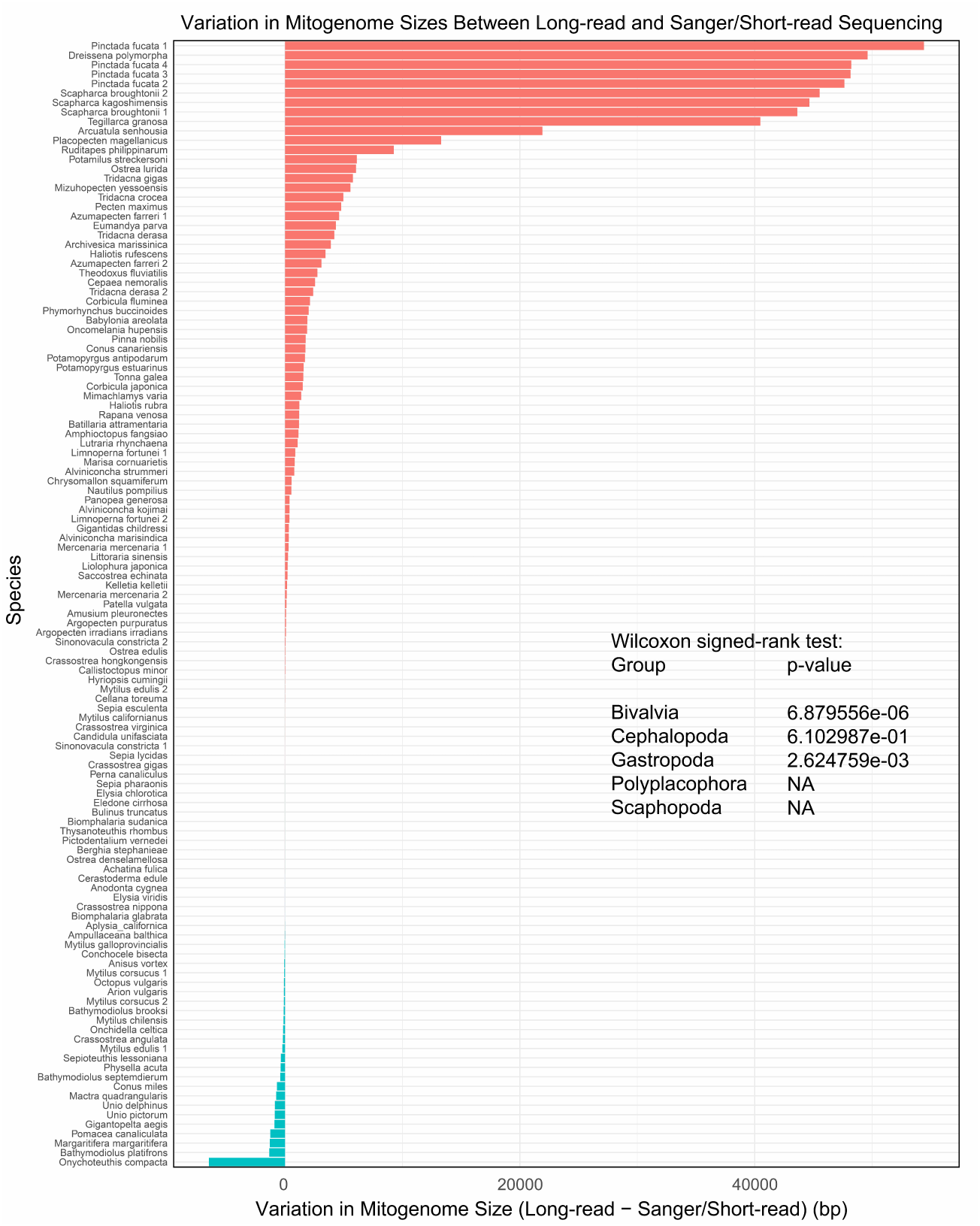

**Fig.S2.**
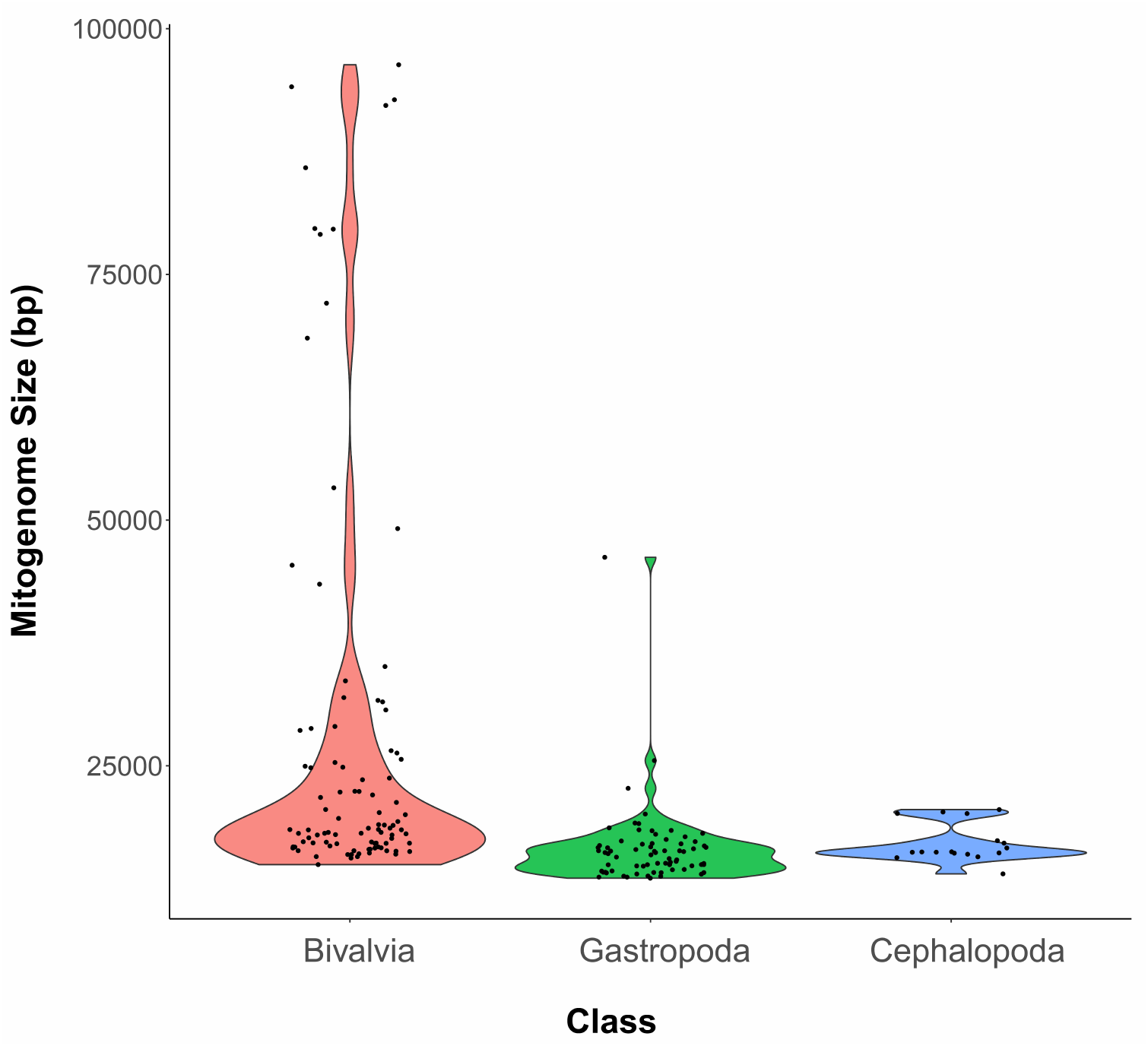

**Fig.S3.**
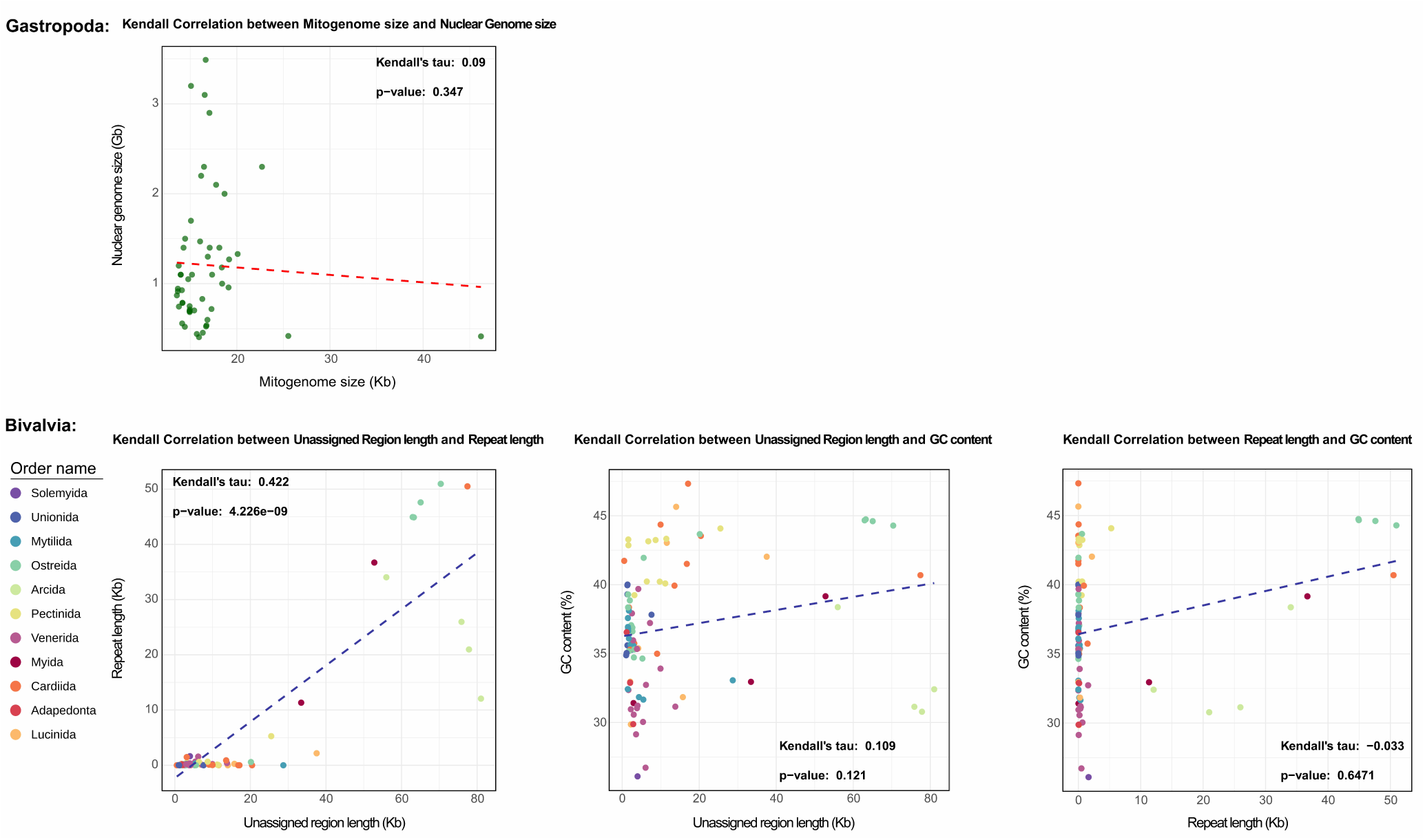

**Fig.S4.**
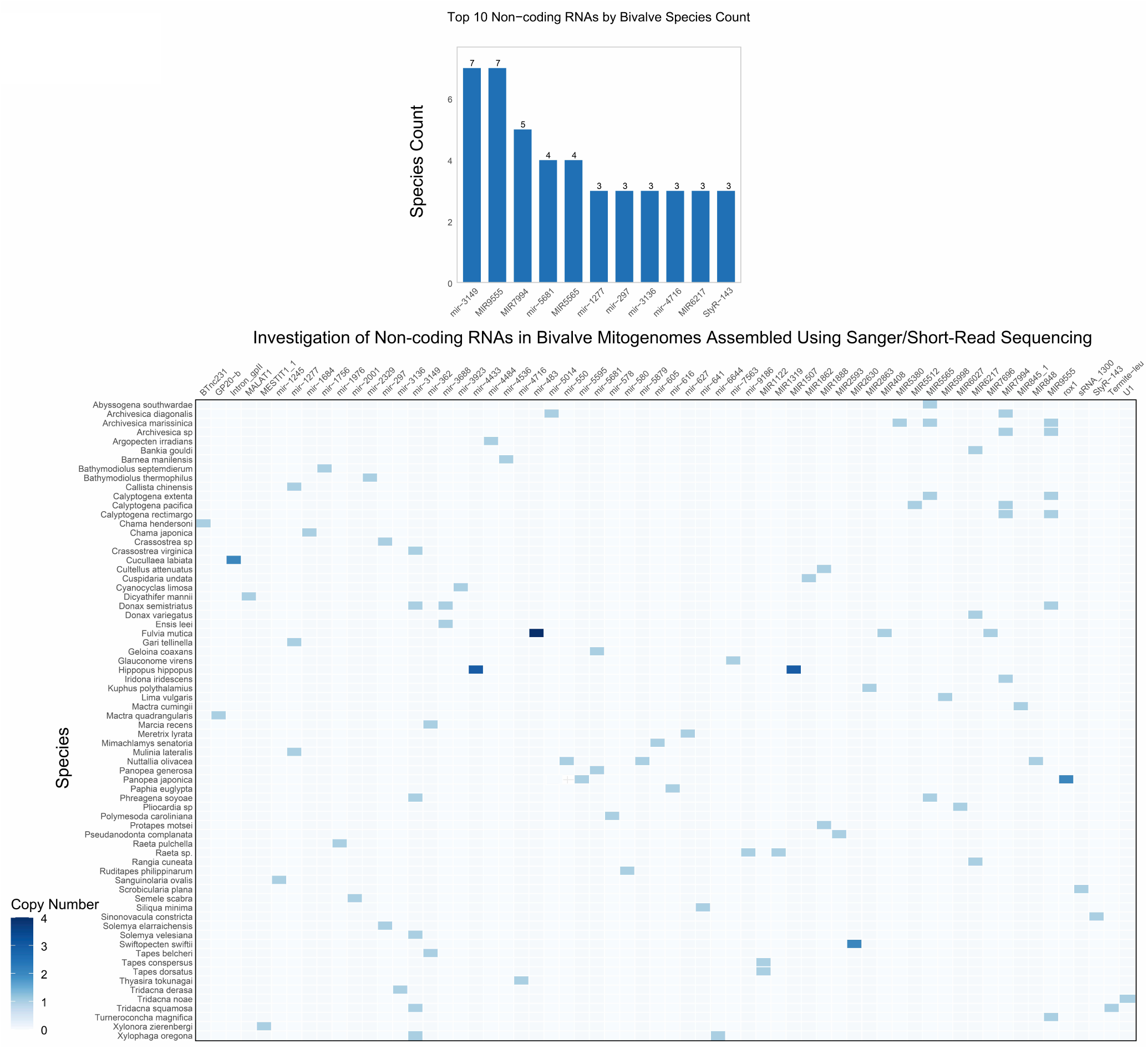

**Fig.S5.**
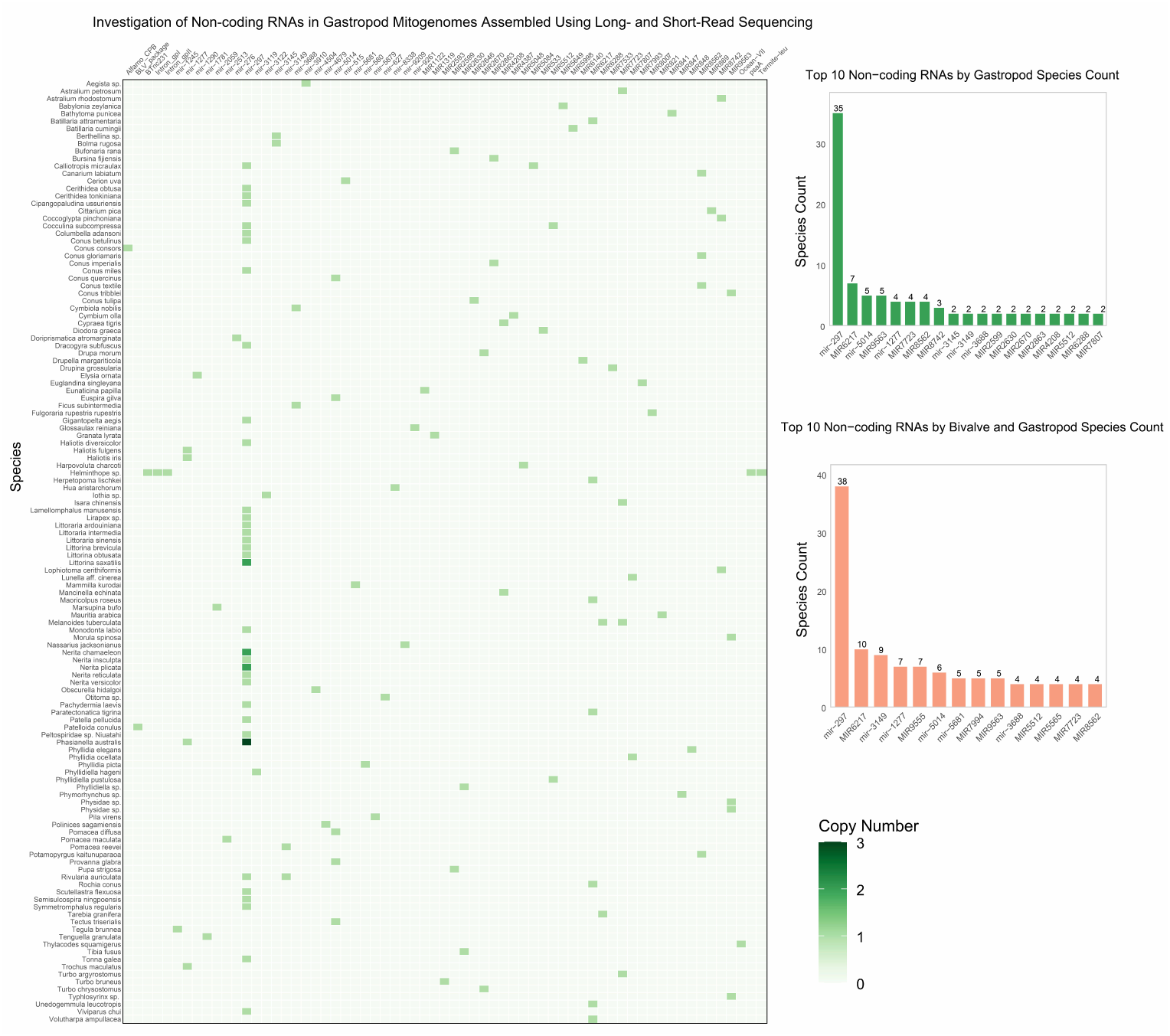

**Fig.S6.**
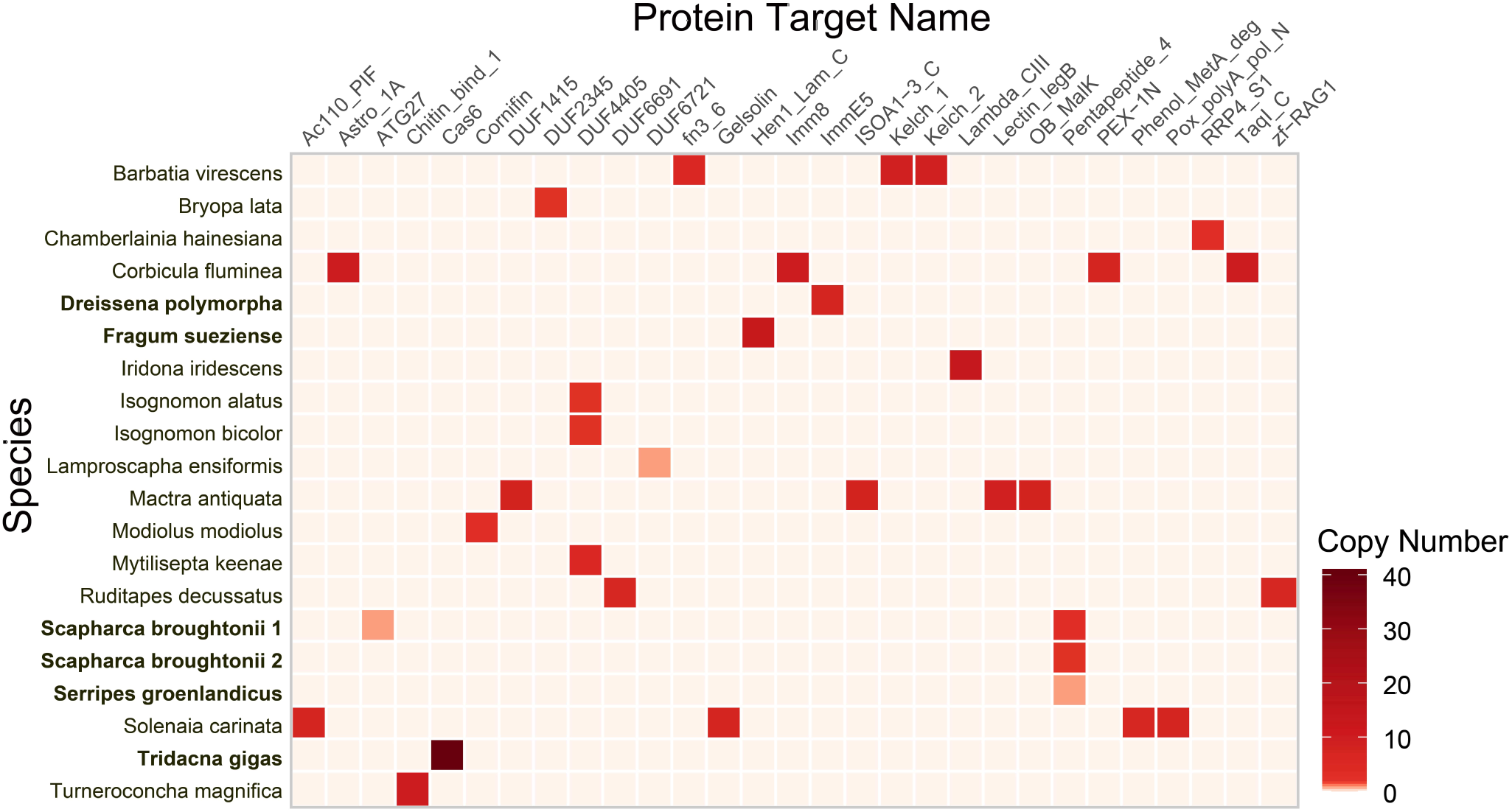

**Fig.S7.**
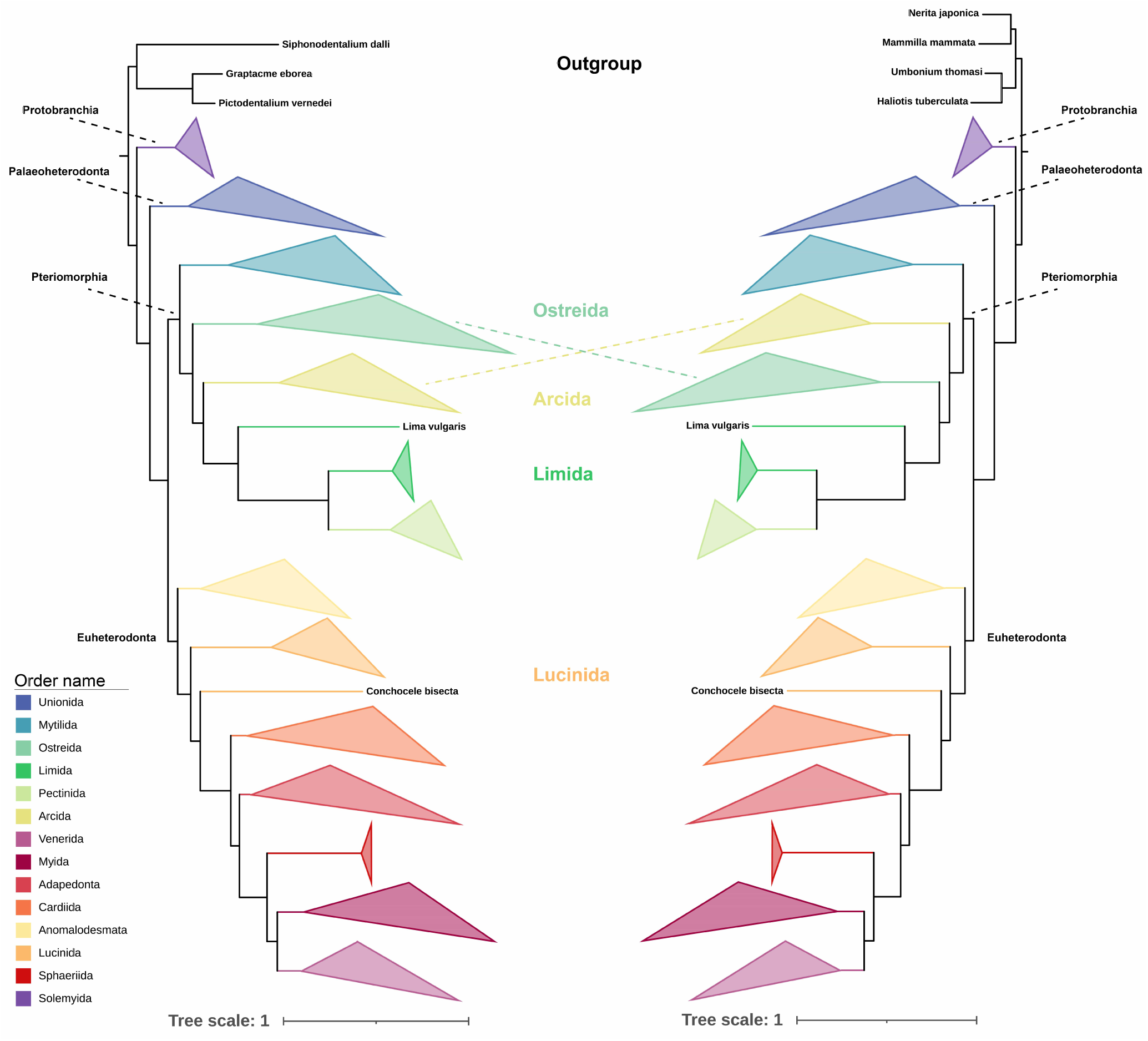

